# The harmonic mean *p*-value for combining dependent tests

**DOI:** 10.1101/171751

**Authors:** Daniel J. Wilson

**Affiliations:** Big Data Institute, Nuffield Department of Medicine, University of Oxford

## Abstract

Analysis of ‘big data’ frequently involves statistical comparison of millions of competing hypotheses to discover hidden processes underlying observed patterns of data, for example in the search for genetic determinants of disease in genome-wide association studies (GWAS). Controlling the family-wise error rate (FWER) is considered the strongest protection against false positives, but makes it difficult to reach the multiple testing-corrected significance threshold. Here I introduce the harmonic mean *p*-value (HMP) which controls the FWER while greatly improving statistical power by combining dependent tests using generalized central limit theorem. I show that the HMP easily combines information to detect statistically significant signals among groups of individually nonsignificant hypotheses in examples of a human GWAS for neuroticism and a joint human-pathogen GWAS for hepatitis C viral load. The HMP simultaneously tests all combinations of hypotheses, allowing the smallest groups of hypotheses that retain significance to be sought. The power of the HMP to detect significant hypothesis groups is greater than the power of the Benjamini-Hochberg procedure to detect significant hypotheses, even though the latter only controls the weaker false discovery rate (FDR). The HMP has broad implications for the analysis of large datasets because it enhances the potential for scientific discovery.

---

Analysis of ‘big data’ has the potential to transform society, not least through improving our understanding of the ways in which genetics influences human traits such as health and disease risk.^1^ However, large datasets present unique challenges. One such challenge now faces geneticists designing future GWAS. To date, participants have typically been typed at around 600,000 genetic variants spread across the 3.2 billion base-pair genome. With the rapidly decreasing costs of DNA sequencing, direct whole genome sequencing (WGS) may soon become routine, raising the possibility of detecting associations at ever more variants.^2,3^ However, this presents a paradox because increasing the number of tests of association requires more stringent *p*-value correction for multiple testing, reducing the probability of detecting any individual association. The idea that analysing more data may lead to fewer discoveries is counter-intuitive, and suggests a flaw of logic.

The problem of testing very many hypotheses while keeping the appropriate false positive rate under control is a longstanding issue in large-scale applications of statistics. The family-wise error rate (FWER) is defined as the probability of falsely rejecting a null in favour of an alternative hypothesis in one or more of all tests performed. Controlling the FWER when some subset of the alternative hypotheses tested might be true is considered the strongest form of protection against false positives.

However, the simple and widely-used Bonferroni method for controlling the FWER tends to be conservative, especially when the individual tests are positively correlated, as often occurs when alternative hypotheses are compared against the same data. In practice, the conservative nature of Bonferroni correction exacerbates the stringent criterion of controlling the FWER, jeopardizing sensitivity to detect true signals.

Alternatives to controlling the FWER have been proposed based on arguments for less stringency. Controlling the false discovery rate (FDR) guarantees that among the significant tests, the proportion in which the null hypothesis is incorrectly rejected in favour of the alternative is limited.^4^ The widely-used Benjamini-Hochberg procedure^4^ for controlling the FDR shares with the Bonferroni method a robustness to positive correlation between individual tests,^5^ but does not share the consequent problem of becoming overly conservative. These advantages have increased the popularity of FDR control, but necessitate the acceptance of a less rigorous standard of control than the FWER, which in practice can produce large numbers of false positives.

Bayesian statistics experiences the same fundamental problem because the posterior odds of any individual hypothesis test are inevitably decreased by increasing the number of alternative hypotheses. However, model averaging using Bayes factors allows alternative hypotheses to be combined, so that comparing a group of alternatives against a common null may rule out the null hypothesis collectively. In the case of GWAS, even if no individual variant shows sufficiently strong evidence of association in a region, the model-averaged signal across that region may still achieve sufficiently strong posterior odds.^6,7^ Combining tests in this way makes an asset of more data by creating the potential for more fine-grained discovery when the signal is sufficiently strong without the liability of requiring that all hypotheses are evaluated individually at the higher level of statistical stringency.

However, there is no general method for combining evidence across hypotheses by model averaging in classical statistics. While some Bayesian arguments advocate simply abandoning classical statistics,^8^ others show that *p*-values from likelihood-based inference are mathematically closely related to Bayesian quantities.^9,10^ Pragmatically, the difficulty of specifying prior information, a tendency for computationally slower methods, and inertia, mean that application of Bayesian methods by practitioners still lags behind classical approaches in many settings, including GWAS. Here I show that competing hypothesis tests can be combined quickly and easily through the harmonic mean *p*-value, improving statistical power and the prospects for discovery using classical statistics, and prompting a reevaluation of the issue of controlling false positive rates in analyses of big data.

## Results

### The harmonic mean *p*-value

For observed data **X** consider *L* mutually exclusive alternative hypotheses *M*_*i*_, *i* = 1 … *L*, all with the same nested null hypothesis *M*_0_. Suppose each alternative has been tested against the null to produce a *p*-value, *p*_*i*_. The main result of this paper is that the weighted harmonic mean *p*-value of any subset *R* of the *p*-values,

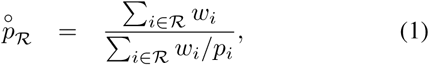

(i) combines the evidence in favour of the group of alternative hypotheses R against the common null, (ii) is an approximately well-calibrated *p*-value for small values, and (iii) the following test controls the strong-sense family-wise error rate (FWER) at level approximately *α* for *α* ≥ 0.05, no matter how many subsets are tested:

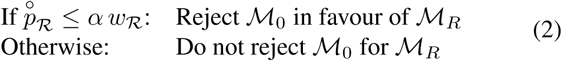

where alternative hypothesis *M*_*i*_ has weight 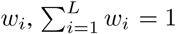 and *w*_*R*_ = Σ_*i*∈*R*_ *w*_*i*_.

Generalized central limit theorem (e.g. ref^11^) can be used to obtain a *p*-value that becomes exact for large groups of hypotheses because 1/p̊_*R*_ tends towards a Landau distribution,^12^ which has probability density function

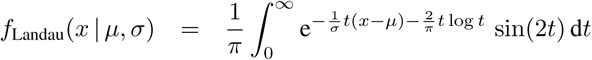

This allows tables of significance thresholds to be computed for interpretation of the HMP (Table 1), and computation of a better-calibrated *p*-value using the HMP as a test statistic:

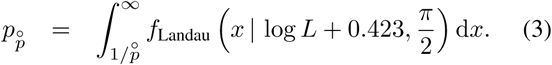

Table 1 shows that direct interpretation of the HMP p̊ tends to be anti-conservative but very closely approximates *p*_p̊_ for small values and small groups of alternative hypotheses.

**Table 1:** Significance thresholds *α*_|*R*|_ for the HMP p̊_*R*_ for varying numbers of alternative hypotheses |*R*| and false positive rates *α*.

| $\vert \mathcal{R}\vert $ | $\alpha = 0.05$ | $\alpha = 0.01$ | $\alpha = 0.001$ |
| --- | --- | --- | --- |
| 10 | 0.040 | 0.0094 | 0.00099 |
| 100 | 0.036 | 0.0092 | 0.00099 |
| 1 000 | 0.034 | 0.0090 | 0.00099 |
| 10 000 | 0.031 | 0.0088 | 0.00098 |
| 100 000 | 0.029 | 0.0086 | 0.00098 |
| 1 000 000 | 0.027 | 0.0084 | 0.00098 |
| 10 000 000 | 0.026 | 0.0083 | 0.00098 |
| 100 000 000 | 0.024 | 0.0081 | 0.00098 |
| 1 000 000 000 | 0.023 | 0.0080 | 0.00097 |

Use of the HMP has several helpful properties that arise from generalized central limit theorem (see Supplementary Methods). It is

1. Robust to positive dependency between the individual *p*-values.
2. Insensitive to the exact number of tests.
3. Robust to the distribution of weights *w*.
4. Most influenced by the smallest *p*-values.

The HMP outperforms Bonferroni and Simes^13^ correction. This latter point means that whenever the Benjamini-Hochberg procedure,^4^ which controls only the FDR, finds significant hypotheses, the HMP will find significant hypotheses or groups of hypotheses. The HMP complements Fisher’s method for combining independent *p*-values,^14^ because the HMP is more appropriate when (i) rejecting the null implies that only one alternative hypothesis may be true, and not all of them (ii) the *p*-values might be positively correlated, and cannot be assumed independent.

In the next section the theory giving rise to the HMP is explained. Readers most interested in application of the HMP can skip to the following sections.

### Model-averaged mean maximum likelihood

A classical analogue of the Bayes factor is the maximized likelihood ratio, which measures the evidence for the alternative hypothesis against the null:

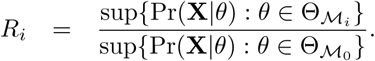

In a likelihood ratio test (LRT), the *p*-value is calculated as the probability of obtaining an *R*_*i*_ as or more extreme if the null hypothesis were true:

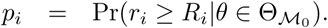

For nested hypotheses (Θ_*M*_0__ ∈ Θ_*M*_*i*__), Wilks’ theorem^15^ approximates the null distribution of *R*_*i*_ as LogGamma(*α* = *V*/2, *β* = 1) when there are *v* degrees of freedom.

The idea motivating this paper was to develop a classical analogue to the model-averaged Bayes factor by deriving the null distribution for the mean maximized likelihood ratio,

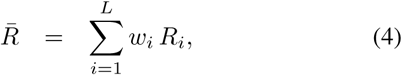

where the weights could take into account prior information and the power of each test. Formally this means the model is treated as a random effect. Choice of weights is considered further in the Supplementary Methods.

The distribution of *R*̄ cannot be approximated by central limit theorem because the LogGamma distribution is heavy tailed, with undefined variance. Instead generalized central limit theorem can be used,^11^ which states that for equal weights (*w*_*i*_ = 1/*L*) and independent and identically distributed *R*_*i*_s,

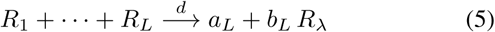

where λ = 1 is the heavy-tail index of the LogGamma(*v*/2,1) distribution, *a*_*L*_ and *b*_*L*_ are constants and *R*_*λ*_ is a Stable distribution with tail index λ. When *v* = 2, the specific form of the Stable distribution is the Landau. The assumptions of equal weights, independence and identical degrees of freedom can be relaxed. Full details of the Stable distribution approximation are in the Supplementary Methods.

Notably, when *v* = 2 and the assumptions of Wilks’ theorem are met, the *p*-value equals the inverse maximized likelihood ratio:

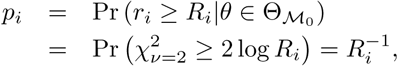

so the mean maximized likelihood ratio equals the inverse HMP:

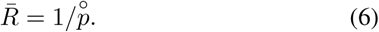

Under these conditions, interpreting *R*̄ and the HMP are exactly equivalent. This equivalence motivates use of the HMP more generally because

1. The HMP will capture similar information to *R*̄ regard-less of the degrees of freedom.
2. The Landau distribution gives an excellent approximation for *R*̄ with *v* = 2, and so for 1/*p*̊.
3. Combining *p*_*i*_s rather than *R*_*i*_s automatically accounts for differences in degrees of freedom.

Further, the HMP is approximately well calibrated because the LogGamma cumulative distribution function is regularly varying, meaning that the model-averaged *p*-value (Equation 3) is approximated by (e.g. ref^16^)

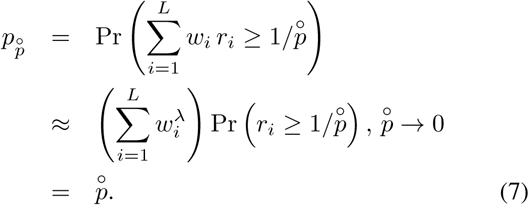

Directly interpreting the HMP using Equation 2 constitutes a multilevel test in the sense that any significant subset of hypotheses implies the HMP of the superset will also be significant because

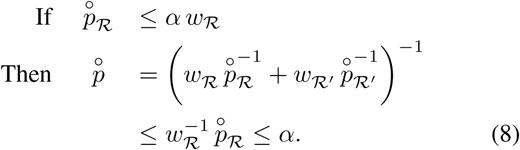

This means that (i) the HMP is a *closed testing procedure*^11^ that controls the strong-sense FWER, (ii) the HMP is more powerful than Bonferroni and Simes correction because the HMP is always smaller than the *p*-values for those tests, and therefore (iii) the HMP will produce significant results whenever the Simes-based Benjamini-Hochberg (BH) procedure does, even though BH only controls the less-stringent FDR. However, these results are only exact when the false positive rate *a* is arbitrarily small. In practice, the exact significance threshold varies by the number of hypotheses combined (Table 1).

So Equation 2 is formally a shortcut procedure that mathematically guarantees *either* superior power over Bonferroni and Simes *or* strong sense control of the FWER depending on whether *α* or *α*_|*R*|_ is employed, respectively. Use of the latter threshold is exact up to the order of the Stable distribution approximation, and equivalent to applying a weighted Bonferroni correction to Equation 3. I recommend the use of this more exact test, available in the R package *harmonicmeanp,* and upon which all subsequent analyses in the main text are based. Analyses based on direct interpretation of the HMP are also presented in the Supplement, and reveal the practical differences between the approaches to be small for *α* = 0.05.

### HMP enables adaptive multiple testing correction by combining *p*-values

That the Bonferroni method for controlling the FWER can be overly stringent, especially when the tests are nonindependent, has long been recognized. In Bonferroni correction, a *p*-value is deemed significant if *p* ≤ *α*/*L*, which becomes more stringent as the number of tests *L* increases. Since human GWAS began routinely testing millions of variants by statistically imputing untyped variants, a new convention was adopted in which a *p*-value is deemed significant if *p* ≤ 5 × 10^‒8^, a rule that implies the effective number of tests is no more than *L* = 10^6^. Several lines of argument were used to justify this *ad hoc* threshold,^19,20^ most applicable only to human GWAS.

In contrast, the HMP affords strong control of the FWER while avoiding both *ad hoc* rules and the undue stringency of Bonferroni correction, an advantage that increases when tests are non-independent. To show how the HMP can recover significant associations among groups of tests that are individually non-significant, I reanalysed a GWAS of neuroticism,^18^ defined as a tendency towards intense or frequent negative emotions and thoughts.^21^ Genotypes were imputed for *L* = 6524432 variants across 170 911 individuals. I used the HMP to perform model-averaged tests of association between neuroticism and variants within contiguous regions of 10, 100 and 1000 kilobases (kb), 10 megabases (Mb), entire chromosomes and the whole genome, assuming equal weights across variants.

Figure 1 shows the *p*-value from Equation 3 for each region *R* adjusted by a factor 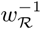 to enable direct comparison to the significance threshold *α* = 0.05. Similar results were obtained from direct interpretation of the HMP (Figure S1). Model averaging tends to make significant and nearsignificant adjusted *p*-values more significant. For example, for every variant significant after Bonferroni correction, the model-averaged *p*-value for the corresponding chromosome was found to be at least as significant.

**Figure 1:**
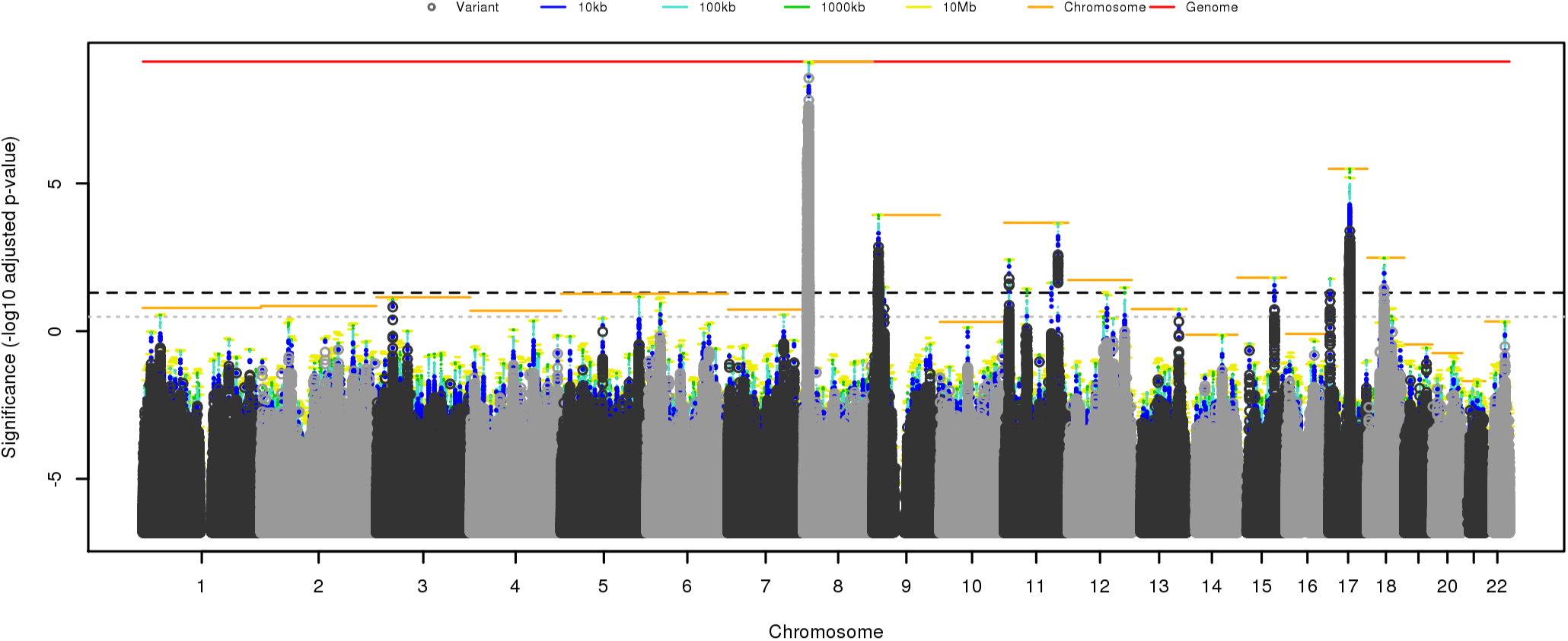
Results of a GWAS of neuroticism in 170911 people.^18^ This Manhattan plot shows the significance of association between neuroticism and *L* = 6524432 variants (dark and light grey points) and overlapping regions of length 10, 100, 1000 and 10 000 kb (blue, cyan, green and yellow bars), entire chromosomes (orange bars) and the whole genome (red bar). Significance is defined as the – log_10_ adjusted *p*-value, where the *p*-value for region *R* is defined by Equation 3, and adjusted by a factor 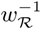 to enable direct comparison to the threshold *α* = 0.05 (black dashed line). The *ad hoc* threshold of *α* = (5 × 10^‒8^)*L* is shown for comparison (grey dotted line).

Model-averaging increases significance more when combining a group of comparably significant *p*-values, e.g. the top hits in chromosome 9. The least improvement is seen when one *p*-value is much more significant than the others, e.g. the top hit in chromosome 3. This behaviour is predicted by the tendency of harmonic means to be dominated by the smallest values. In the extreme case that one *p*-value dominates the significance of all others, the HMP test becomes equivalent to Bonferroni correction. This implies that Bonferroni correction might not be improved upon for ‘needle-in-a-haystack’ problems. Conversely, dependency among tests actually *improves* the sensitivity of the HMP because one significant test may be accompanied by other correlated tests that collectively reduce the harmonic mean *p*-value.

In some cases, the HMP found significant regions where none of the individual variants were significant. For example, no variants on chromosome 12 were significant by Bonferroni correction nor by the *ad hoc* genome-wide significance threshold of 5 × 10^‒8^. However, the HMP found significant 10Mb regions spanning several peaks of non-significant individual *p*-values. One of those, variant rs7973260, which showed an individual *p*-value for association with neuroticism of 2.4 × 10^‒7^, had been reported as also associated with depressive symptoms (*p* =1.8 × 10^‒9^). Such cross-association or ‘quasi-replication’, in which a variant is near-significant for the trait-of-interest and significant for a related trait, can be regarded as providing additional support for the variant’s involvement in the trait-of-interest.^18^

In chromosome 3, individual variants were found to be significant by the *ad hoc* threshold of 5 × 10^‒8^, but neither Bonferroni correction nor the HMP agreed those variants or regions were significant at a FWER of *×* = 0.05. Indeed the HMP found chromosome 3 non-significant as a whole. Variant rs35688236, which had the smallest *p*-value on chromosome 3 of 2.4 × 10^‒8^, had not validated when tested in a quasi-replication exercise that involved testing variants associated with neuroticism for association with subjective wellbeing or depressive symptoms.^18^

These observations illustrate that the HMP adaptively combines information among groups of similarly significant tests where possible, while leaving lone significant tests subject to Bonferroni-like stringency, providing a general approach to combining *p*-values that does not require specific knowledge of the dependency structure between tests.

### HMP allows large-scale testing for higher-order interactions without punitive thresholds

Scientific discovery is currently hindered by avoidance of large-scale exploratory hypothesis testing for fear of attracting multiple testing correction thresholds that render signals found by more limited testing no longer significant. A good example is the approach to testing for pairwise or higher-order interactions between variants in GWAS. The Bonferroni threshold for testing all pairwise interactions invites a threshold (*L* + 1)/2 times more stringent than the threshold for testing variants individually, and strictly speaking this must be applied to *every* test, even though this is highly conservative because of the dependency between tests. The alternative of controlling the FDR risks a high probability of falsely detecting artefacts among any genuine associations discovered. Therefore interactions are not usually tested for.

To show how model-averaging using the HMP greatly alleviates this problem, I reanalysed human and pathogen genetic variants from a GWAS of pre-treatment viral load in hepatitis C virus (HCV)-infected patients.^22^ Jointly analysing the influence of human and pathogen variation on infection is an area of great interest, but requires a Bonferroni threshold of *α*/(*L*_*H*_ + *L*_*P*_) when there are *L*_*H*_ and *L*_*P*_ variants in the human and pathogen genomes respectively, compared to *α*/(*L*_*H*_ + *L*_*P*_) if testing the human and pathogen variants separately. In this example, *L*_*H*_ = 399 420 and *L*_*P*_ = 827.

In the original study, a known association with viral load was replicated at human chromosome 19 variant rs12979860 in *IFNL4* (*p* = 5.9 × 10^‒10^), below the Bonferroni threshold of 1.3 × 10^‒7^. The most significant pairwise interaction I found, assuming equal weights, involved the adjacent variant, rs8099917, with *p* = 2.2 × 10^‒10^. However, this did not meet the more stringent Bonferroni threshold of 1.5 × 10^‒10^ (Figure 2A). If the original study’s authors had performed and reported all 330 million tests, they could have been compelled to declare the marginal association in *IFNL4* non-significant, despite what intuitively appears like a clear signal.

**Figure 2:**
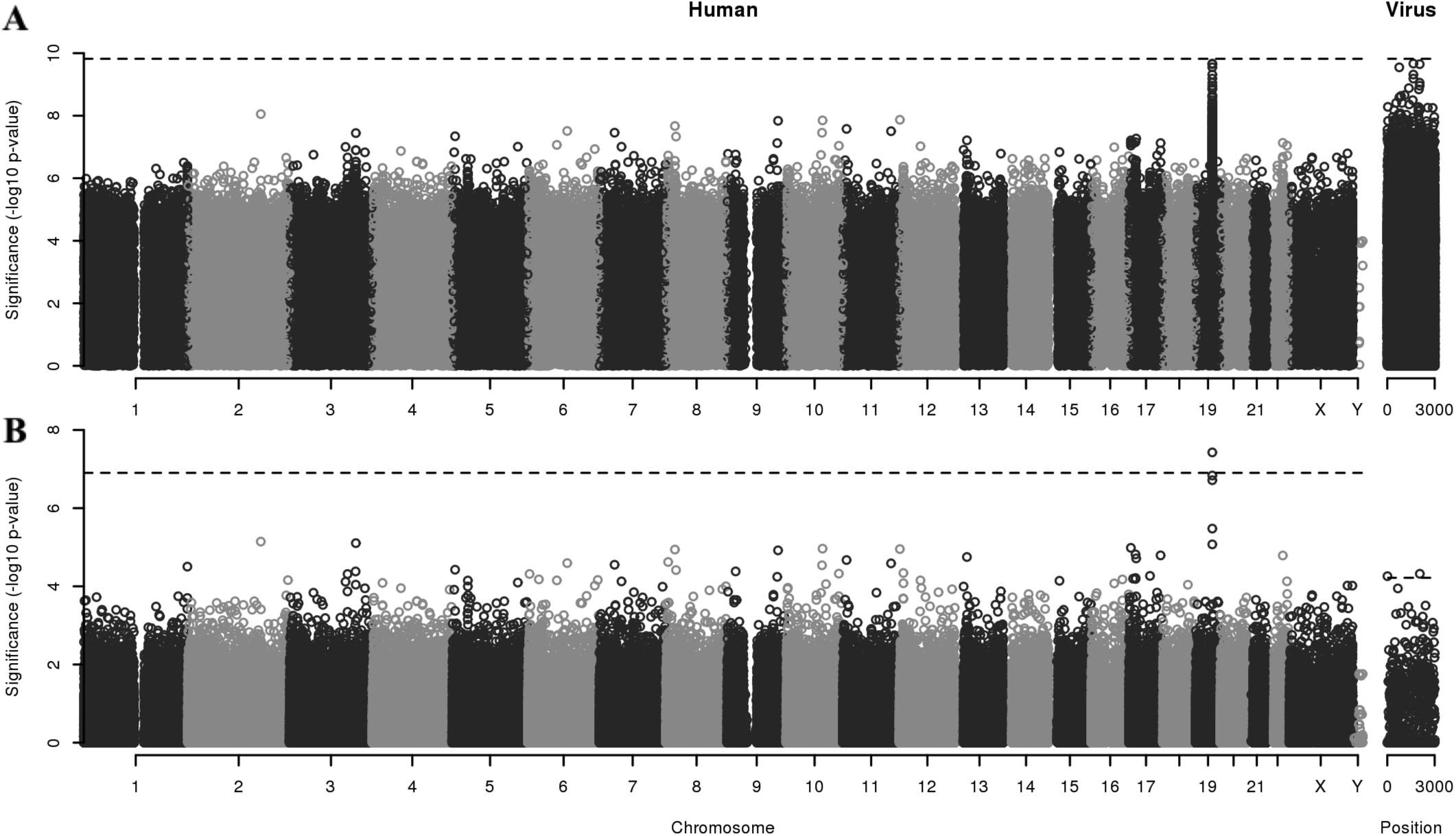
Joint human-pathogen GWAS reanalysis of viral load in 410 HCV genotype 3a-infected white Europeans.^22^ All pairs of human nucleotide variants and viral amino acid variants were tested for association. Interactions between human and virus variants’ effects on viral load were not constrained to be additive. **A**. Significance of 330 320 340 tests plotted by position of both the human and viral variant. **B**. Significance of 399 420 human variants model-averaged using the HMP over every possible interaction with 827 viral variants and *vice versa*. The significance thresholds controlling the FWER at *α* = 0.05 are indicated (black dashed lines): *α*/(*L*_*H*_ + *L*_*p*_), *α*/*L*_*H*_ and *α*/*L*_*p*_.

Model averaging using the HMP reduces this disincentive to perform additional related tests. Figure 2B shows that despite no significant pairwise tests involving rs8099917, model averaging recovered a combined *p*-value of 3.7 × 10^‒8^, below the multiple testing threshold of 1.3 × 10^‒7^. Additionally, two viral variants produced statistically significant model-averaged *p*-values of 5.5 × 10^‒5^ and 4.8 × 10^‒5^ at polyprotein positions 10 and 2 061 in the capsid and NS5a zinc finger domain (GenBank AQW44528), below the multiple testing threshold of 6.0 × 10^‒5^.

These results show how model-averaging using the HMP can enhance scientific discovery by (i) encouraging tests for higher order interactions when they otherwise would not be attempted and (ii) recovering lost signals of marginal associations after performing an ‘excessive’ number of tests.

### Untangling the signals driving significant model-averaged *p*-values

When more than one alternative hypothesis is found to be significant, either individually or as part of a group, it is desirable to quantify the relative strength of evidence in favour of the competing alternatives. This is particularly true when disentangling the contributions of a group of individually nonsignificant alternatives that are significant only in combination.

Sellke, Bayarri and Berger^9^ proposed a conversion from *p*-values into Bayes factors which, when combined with prior information and test power through the model weights, produces posterior model probabilities and credible sets of alternative hypotheses. The Supplementary Methods detail how the Bayes factors are approximately proportional to the weighted inverse *p*-value. This linearity mirrors the HMP itself, whose inverse is an arithmetic mean of the inverse *p*-values.

After conditioning on rejection of the null hypothesis by normalizing the approximate model probabilities to sum to 100%, the probability that the association involved human variant rs8099917 was 54.4%. This signal was driven primarily by the three viral variants with the highest probability of interacting with rs8099917 in their effect on pre-treatment viral load: position 10 in the capsid (10.9%), position 669 in the E2 envelope (8.7%) and position 2061 in the NS5a zinc finger domain (11.4%) (Figure 3). Even though the model-averaged *p*-value for the envelope variant was not itself significant, this revealed a plausible interaction between it and the most significant human variant rs8099917.

**Figure 3:**
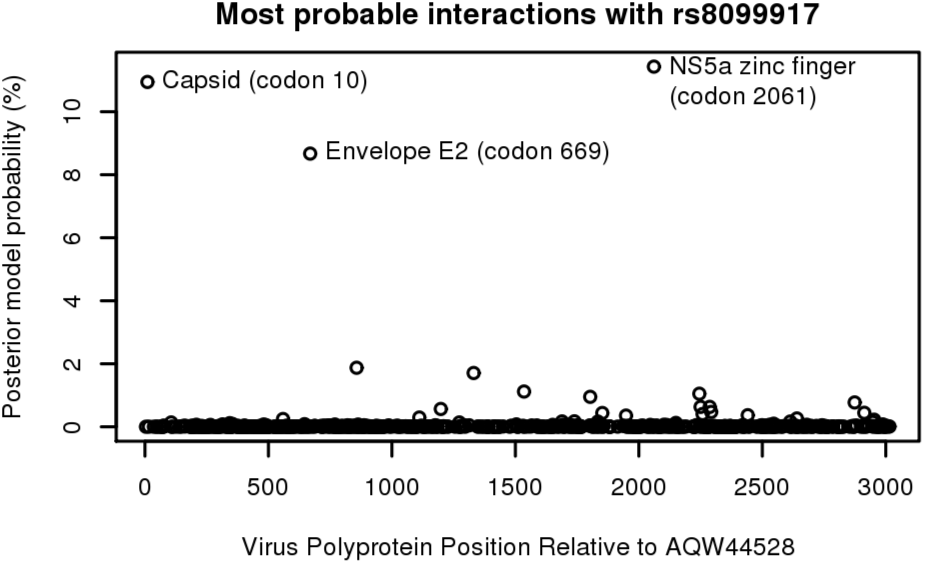
In the joint human-HCV GWAS, the approximate posterior probability of association with rs8099917 was 54.4% in total, with the most probable interactions involving three polyprotein positions.

## Discussion

The HMP provides a way to calculate model-averaged *p*-values, providing a powerful and general method for combining tests while controlling the strong-sense FWER. It provides an alternative to both the overly conservative Bonferroni control of the FWER, and the lower stringency of FDR control. The HMP allows the incorporation of prior information through model weights, and is robust to positive dependency between the *p*-values. The HMP is approximately well-calibrated for small values, while a null distribution, derived from generalized central limit theorem, is easily computed. When the HMP is not significant, neither is any subset of the constituent tests.

The HMP is more appropriate for combining *p*-values than Fisher’s method when the alternative hypotheses are mutually exclusive, as in model comparison. When the alternative hypotheses all have the same nested null hypothesis, the HMP is interpreted in terms of a model-averaged likelihood ratio test. However, the HMP can be used more generally to combine tests that are not necessarily mutually exclusive, but which may have positive dependency. It can be used alone or in combination, for example with Fisher’s method to combine model-averaged *p*-values between groups of independent data.

The theory underlying the HMP provides a fundamentally different way to think about controlling the FWER through multiple testing correction. The Bonferroni threshold increases linearly with the number of tests, whereas the HMP is the reciprocal of the mean inverse *p*-value. To maintain significance with Bonferroni correction, the minimum *p*-value must decrease linearly as the number of tests increases. This strongly penalizes exploratory analyses. In contrast, when the false positive rate *a* is small, to maintain significance with the HMP requires only that the mean inverse *p*-value remains constant as the number of tests increases. This does not penalize exploratory analyses so long as the ‘quality’ of the additional hypotheses tested, measured by the inverse *p*-value, does not decline.

Through example applications to GWAS, I have shown that the HMP combines tests adaptively, producing Bonferroni-like adjusted *p*-values for ‘needle-in-a-haystack’ problems when one test dominates, but able to capitalize on numerous strongly significant tests to produce smaller adjusted *p*-values when warranted. I have shown how model averaging using the HMP encourages exploratory analysis and can recover signals of significance among groups of individually non-significant tests, properties that have the potential to enhance the scientific discovery process.

## Acknowledgements

DJW is a Sir Henry Dale Fellow, jointly funded by the Wellcome Trust and the Royal Society (Grant 101237/Z/13/Z) and a member of the STOP-HCV consortium, which is funded by an award from the Medical Research Council (MR/K01532X/1). I thank the Social Science Genetic Association Consortium, the STOP-HCV Consortium and HCV Research UK Biobank for sharing data, Azim Ansari, Vincent Pedergnana and Chris Spencer for sharing expertize and Simon Myers for helpful comments.

## Software Availability

An R package implementing the harmonic mean *p*-value and MAMML tests is available from https://cran.r-project.org/package=harmonicmeanp.

## References

[1] The Royal Society. Machine learning: the power and promise of computers that learn by example, 2017.

[2] Christian Fuchsberger, Jason Flannick, Tanya M Teslovich, Anubha Mahajan, Vineeta Agarwala, Kyle J Gaulton, Clement Ma, Pierre Fontanillas, Loukas Moutsianas, Davis J McCarthy, et al. The genetic architecture of type 2 diabetes. Nature, 536(7614):41–47, 2016.

[3] Wouter van Rheenen, Aleksey Shatunov, Annelot M Dekker, Russell L McLaughlin, Frank P Diekstra, Sara L Pulit, Rick AA van der Spek, Urmo Vosa, Simone de Jong, Matthew R Robinson, et al. Genome-wide association analyses identify new risk variants and the genetic architecture of amyotrophic lateral sclerosis. Nature genetics, 48(9):1043–1048, 2016.

[4] Yoav Benjamini and Yosef Hochberg. Controlling the false discovery rate: a practical and powerful approach to multiple testing. Journal of the royal statistical society. Series B (Methodological), pages 289–300, 1995.

[5] Yoav Benjamini and Daniel Yekutieli. The control of the false discovery rate in multiple testing under dependency. Annals of statistics, pages 1165–1188, 2001.

[6] Teresa Ferreira and Jonathan Marchini. Modeling interactions with known risk loci: a bayesian model averaging approach. Annals of human genetics, 75(1):1–9, 2011.

[7] Lorin Crawford, Ping Zeng, Sayan Mukherjee, and Xiang Zhou. Detecting epistasis with the marginal epistasis test in genetic mapping studies of quantitative traits. PLoS Genetics, 13(7):e1006869, 2017.

[8] James O Berger and Thomas Sellke. Testing a point null hypothesis: The irreconcilability of p values and evidence. Journal of the American statistical Association, 82(397):112–122, 1987.

[9] Thomas Sellke, M. J. Bayarri, and James O. Berger. Calibration of p values for testing precise null hypotheses. The American Statistician, 55(1):62–71, 2001.

[10] Kenneth Rice. A decision-theoretic formulation of fisher?s approach to testing. The American Statistician, 64(4):345–349, 2010.

[11] Vladimir V Uchaikin and Vladimir M Zolotarev. Chance and stability: stable distributions and their applications. Walter de Gruyter, 1999.

[12] Lev Davidovich Landau. On the energy loss of fast particles by ionization. J. Phys., 8:201–205, 1944.

[13] R John Simes. An improved bonferroni procedure for multiple tests of significance. Biometrika, 73(3):751–754, 1986.

[14] Ronald Aylmer Fisher. Statistical Methods for Research Workers. Oliver and Boyd, Edinburgh, fifth edition, 1934.

[15] Samuel S Wilks. The large-sample distribution of the likelihood ratio for testing composite hypotheses. The Annals of Mathematical Statistics, 9(1):60–62, 1938.

[16] Thomas Mikosch. Regular variation, subexponentiality and their applications in probability theory. Eindhoven University of Technology, 1999.

[17] Ruth Marcus, Peritz Eric, and K Ruben Gabriel. On closed testing procedures with special reference to ordered analysis of variance. Biometrika, 63(3):655–660, 1976.

[18] Aysu Okbay, Bart ML Baselmans, Jan-Emmanuel De Neve, Patrick Turley, Michel G Nivard, Mark Alan Fontana, S Fleur W Meddens, Richard Karlsson Linner, Cornelius A Rietveld, Jaime Derringer, et al. Genetic variants associated with subjective well-being, depressive symptoms, and neuroticism identified through genome-wide analyses. Nature genetics, 48(6):624–633, 2016.

[19] Itsik Pe’er, Roman Yelensky, David Altshuler, and Mark J Daly. Estimation of the multiple testing burden for genomewide association studies of nearly all common variants. Genetic epidemiology, 32(4):381–385, 2008.

[20] Joao Fadista, Alisa K Manning, Jose C Florez, and Leif Groop. The (in) famous gwas p-value threshold revisited and updated for low-frequency variants. European Journal of Human Genetics, 24(8):1202–1205, 2016.

[21] Robert R McCrae and Paul T Costa Jr. The five-factor theory of personality. In L.A. Pervin O.P. John, R.W. Robins, editor, Handbook of personality: Theory and research, pages 159–181. Guilford Press, New York, third edition, 2008.

[22] M Azim Ansari, Vincent Pedergnana, Camilla LC Ip, Andrea Magri, Annette Von Delft, David Bonsall, Nimisha Chaturvedi, Istvan Bartha, David Smith, George Nicholson, et al. Genome-to-genome analysis highlights the effect of the human innate and adaptive immune systems on the hepatitis c virus. Nature Genetics, 49(5):666–673, 2017.

[23] Jie Zheng, A Mesut Erzurumluoglu, Benjamin L Elsworth, John P Kemp, Laurence Howe, Philip C Haycock, Gibran Hemani, Katherine Tansey, Charles Laurin, Beate St Pourcain, et al. Ld hub: a centralized database and web interface to perform ld score regression that maximizes the potential of summary level gwas data for snp heritability and genetic correlation analysis. Bioinformatics, 33(2):272–279, 2017.

[24] Cristen J Willer, Yun Li, and Goncalo R Abecasis. Metal: fast and efficient meta-analysis of genomewide association scans. Bioinformatics, 26(17):2190–2191, 2010.

[25] Xavier Didelot and Daniel J Wilson. Clonal-frameml: efficient inference of recombination in whole bacterial genomes. PLoS computational biology, 11(2):e1004041, 2015.

[26] Stéphane Guindon, Jean-François Dufayard, Vincent Lefort, Maria Anisimova, Wim Hordijk, and Olivier Gascuel. New algorithms and methods to estimate maximum-likelihood phylogenies: assessing the performance of phyml 3.0. Systematic biology, 59(3):307–321, 2010.

[27] Jovan Karamata. Sur un mode de croissance régulière. théorèmes fondamentaux. Bull. Soc. Math. France, 61:55–62, 1933.

[28] IV Zaliapin, Yan Y Kagan, and Federic P Schoenberg. Approximating the distribution of pareto sums. Pure and Applied geophysics, 162(6):1187–1228, 2005.

[29] Robert H Rimmer and John P Nolan. Stable distributions in mathematica. Mathematica Journal, 9(4):776–789, 2005.

[30] Geoff Robinson. FMStable: Finite Moment Stable Distributions, 2012. R package version 0.1-2.

[31] Katarzyna Bartkiewicz, Adam Jakubowski, Thomas Mikosch, and Olivier Wintenberger. Stable limits for sums of dependent infinite variance random variables. Probability Theory and Related Fields, 150(3):337–372, 2011.

[32] Yosef Hochberg and Uri Liberman. An extended simes’ test. Statistics & Probability Letters, 21(2):101–105, 1994.

[33] Aaditya Ramdas, Rina Foygel Barber, Martin J Wainwright, and Michael I Jordan. A unified treatment of multiple testing with prior knowledge. arXiv preprint arXiv:1703.06222, 2017.

[34] Thomas M Loughin. A systematic comparison of methods for combining p-values from independent tests. Computational statistics & data analysis, 47(3):467–485, 2004.

[35] HO Lancaster. The combination of probabilities: an application of orthonormal functions. Australian & New Zealand Journal of Statistics, 3(1):20–33, 1961.

[36] T Liptak. On the combination of independent tests. Magyar Tud Akad Mat Kutato Int Kozl, 3:171–197, 1958.

[37] Eugene S Edgington. An additive method for combining probability values from independent experiments. The Journal of Psychology, 80(2):351–363, 1972.

[38] G S Mudholkar and EO George. The logic method for combining probabilities. In J Rustagi, editor, Symposium on Optimizing Methods in Statistics, pages 345–366. Academic Press, New York, 1979.

[39] Gideon Schwarz et al. Estimating the dimension of a model. The annals of statistics, 6(2):461–464, 1978.

[40] Adrian E Raftery. Approximate bayes factors and accounting for model uncertainty in generalised linear models. Biometrika, 83(2):251–266, 1996.

[41] Doug Speed, Na Cai, Michael R Johnson, Sergey Nejentsev, David J Balding, UCLEB Consortium, et al. Reevaluation of snp heritability in complex human traits. Nature Genetics, 49:986–992, 2017.

[42] Matthew Stephens and David J Balding. Bayesian statistical methods for genetic association studies. Nature reviews. Genetics, 10(10):681, 2009.

[43] Bradley Efron, Robert Tibshirani, John D Storey, and Virginia Tusher. Empirical bayes analysis of a microarray experiment. Journal of the American statistical association, 96(456):1151–1160, 2001.

[44] John D Storey and Robert Tibshirani. Statistical significance for genomewide studies. Proceedings of the National Academy of Sciences, 100(16):9440–9445, 2003.

[45] John D Storey. A direct approach to false discovery rates. Journal of the Royal Statistical Society: Series B (Statistical Methodology), 64(3):479–498, 2002.

[46] Matthew Stephens. False discovery rates: a new deal. Biostatistics, 18(2):275–294, 2016.

[47] Claire Chewapreecha, Pekka Marttinen, Nicholas J Croucher, Susannah J Salter, Simon R Harris, Alison E Mather, William P Hanage, David Goldblatt, Francois H Nosten, Claudia Turner, et al. Comprehensive identification of single nucleotide polymorphisms associated with beta-lactam resistance within pneumococcal mosaic genes. PLoS genetics, 10(8):e1004547, 2014.

[48] Sarah G Earle, Chieh-Hsi Wu, Jane Charlesworth, Nicole Stoesser, N Claire Gordon, Timothy M Walker, Chris CA Spencer, Zamin Iqbal, David A Clifton, Katie L Hopkins, et al. Identifying lineage effects when controlling for population structure improves power in bacterial association studies. Nature microbiology, 1:16041,2016.

